# *Onecut1* partially contributes to the liver progenitor cell transition and acquisition of metastatic potential in hepatocellular carcinoma

**DOI:** 10.1101/2022.09.20.508738

**Authors:** Yu Liu, Hiroaki Higashitsuji, Katsuhiko Itoh, Kanji Yamaguchi, Atsushi Umemura, Yoshito Itoh, Jun Fujita

**Affiliations:** Department of Clinical Molecular Biology, Graduate School of Medicine, Kyoto University, Kyoto 606-8507, Japan.; Department of Gastroenterology and Hepatology, Kyoto Prefectural University of Medicine, Kyoto 602-8566, Japan.; Educational Development Center, Kyoto University of Advanced Science, Kyoto 615-8577, Japan.; Department of Pharmacology, Kyoto Prefectural University of Medicine, Kyoto 602-8566, Japan.; Department of Molecular Genetics, Graduate School of Medicine, Kyoto University, Kyoto 606-8507, Japan.

## Abstract

Metastasis-initiating cells are considered to originate from stem cell-like cancer cells. In hepatocellular carcinoma, liver progenitor-like cells are reported to be derived from hepatocytes, indicating the possible acquisition of metastatic potential during hepatocyte-to-cholangiocyte transdifferentiation. Consistent with the expression pattern observed during ductal plate formation, we revealed an LPC transition with *Onecut1* accumulation both during hepatocyte-to-cholangiocyte transdifferentiation and in a cell model. This event may be associated with transient acquisition of metastatic potential.

## Introduction

In hepatocellular carcinoma (HCC), expression of cholangiocyte/liver progenitor cell (LPC) markers (i.e., *Krt19*, *Epcam*, *Sox9* and *Spp1*) indicates poor prognosis (Govaere et al. 2014; Chaudhary et al. 2018; Sun et al. 2013; Guo et al. 2012; Pan et al. 2003). However, the origin of LPCs in HCC, as well as the origin of HCC itself, have been debated, and the evidence is conflicting. In 2015, Mu *et al*. (Mu et al. 2015) reported that in both genotoxic and genetic models, HCC cells, including KRT19^+^ cells within tumors, arose exclusively from hepatocytes, indicating a crucial role of hepatocyte-to-cholangiocyte transdifferentiation in stemness acquisition.

*One cut homeobox 1* (*Onecut1*; also known as *Hnf6*) is a key regulator during the development but not the postnatal regeneration of the liver and biliary system (Clotman et al. 2002; Walter et al. 2014; Schaub et al. 2018). In the fetal liver, ONECUT1 is weakly expressed in the liver parenchyma and accumulates in the ductal plate. In the adult liver, ONECUT1 is expressed in hepatocytes, and its expression is completely lost in mature cholangiocytes (Limaye et al. 2008). *Onecut1* knockout mice show absence of the gallbladder and interruption of the bile duct with expansion of prematurely differentiated cholangiocytes, suggesting that *Onecut1* inhibits premature cholangiocyte differentiation.

## Results

### ONECUT1 facilitates incomplete hepatocyte-to-cholangiocyte transdifferentiation

Two cell lines, hepaEP1 and hepaEP2 (hepaEPs), with a more epithelial-like morphology than the parental cells, were subcloned from the mouse HCC cell line Hepa1-6 by limiting dilution (Figure 1A). In hepaEPs, the mRNA level of *Alb* was significantly increased, indicating hepatic differentiation (Figure 1B). Elevated ALB secretion was verified by immunoblotting of serum-free culture supernatants and immunoprecipitates from serum-containing culture supernatants (Figure 1C, S1A). Proliferation of hepaEP1 slightly increased *in vitro* but not in xenograft models (Figure S1B, S1C).

**Figure 1.**
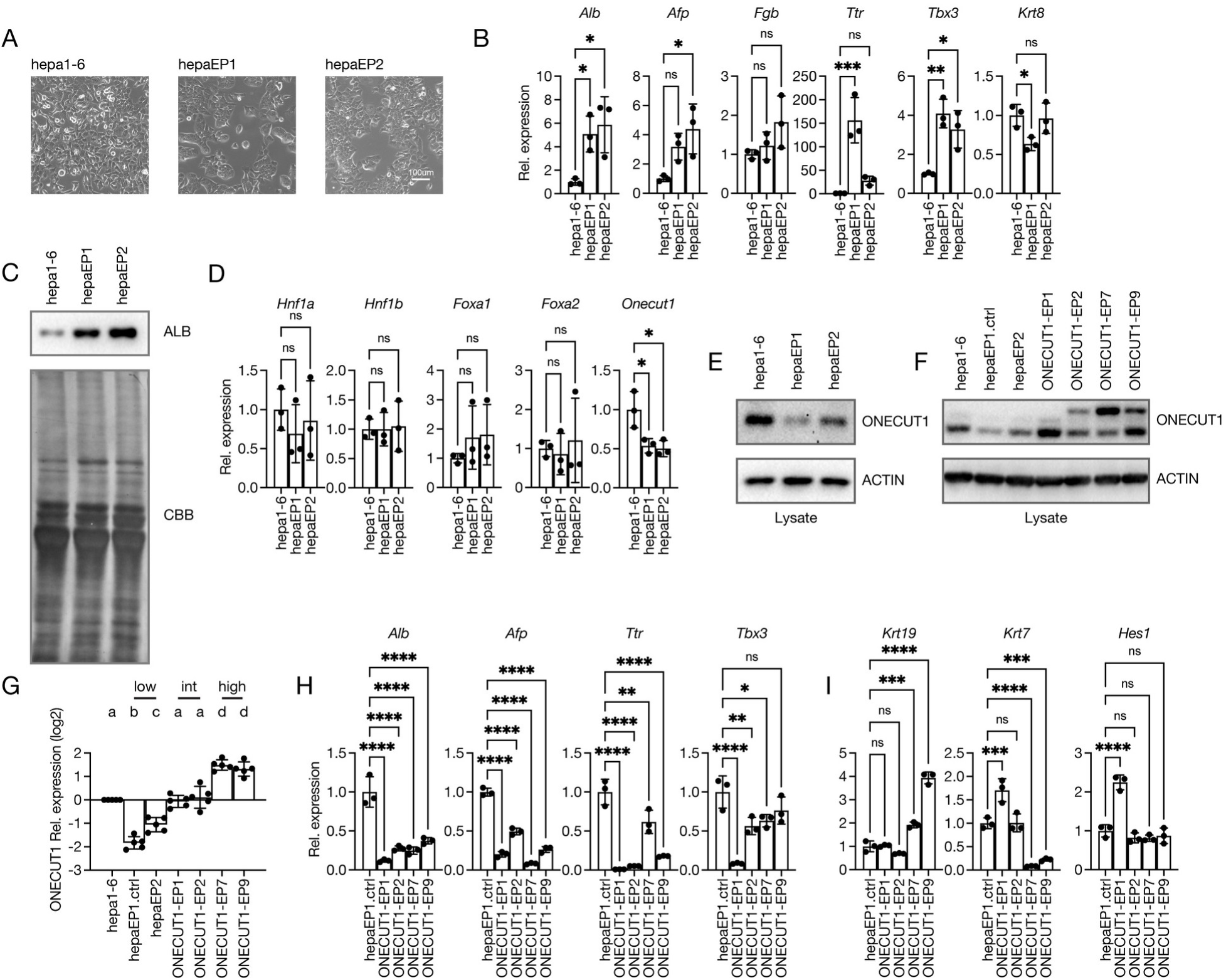
Gradual expression of ONECUT1 in ONECUT1^low^ cells facilitates incomplete hepatocyte-to-cholangiocyte transdifferentiation. **(A)** Morphology of the mouse HCC cell line Hepa1-6 and the epithelial-like hepaEP1 and hepaEP2 cell lines derived from Hepa1-6 cells. **(B)** Expression of hepatic markers, as measured by quantitative RT‒PCR. The mRNA level of Hepa1-6 was set to 1; n = 3; * indicates a significant difference by 1-way ANOVA. **(C)** Immunoblotting of secreted ALB (upper) and total protein (lower). **(D)** Expression of liver-enriched transcription factors, as measured by quantitative RT‒PCR. The mRNA level of Hepa1-6 was set to 1; n = 3; * indicates a significant difference by 1-way ANOVA. **(E, F)** Immunoblotting of ONECUT1 (upper) and ACTIN (lower). **(G)** Quantification of the protein levels in **(F)**. The protein level of Hepa1-6 was set to 1; n = 5; the different letters indicate significant differences by 1-way ANOVA. **(H, I)** Expression of hepatic or cholangiocyte markers, as measured by quantitative RT‒PCR. The mRNA level of hepaEP1 was set to 1; n = 3; * indicates a significant difference by 1-way ANOVA.

To investigate whether liver-enriched transcription factors play a role in the hepatic differentiation of Hepa1-6 cells, we analyzed the mRNA expression of *Hnf1a*, *Hnf1b*, *Foxa1*, *Foxa2*, *Foxa3*, *Hnf4a*, *Onecut1*, *Cebpa* and *Cebpb*. *Foxa3*, *Hnf4a*, *Cebpa* and *Cebpb* were not detected, while the mRNA expression of only *Onecut1* showed a significant decrease in hepaEPs (Figure 1D). The decrease in ONECUT1 protein expression in hepaEPs was verified by immunoblotting of cell lysates (Figure 1E).

To understand the role of *Onecut1*, we generated stable clones by introducing plasmids expressing ONECUT1 or 3FLAG-ONECUT1 into hepatically differentiated hepaEP1 cells (Figure 1F). The obtained cell lines and hepaEPs (hereafter collectively referred to as ONECUT1-EPs) were divided into 3 groups: ONECUT1^low^-EPs (hepaEP1 and hepaEP2), ONECUT1^int^-EPs (ONECUT1-EP1 and ONECUT1-EP2) and ONECUT1^high^-EPs (ONECUT1-EP7 and ONECUT1-EP9) with more than 2-fold lower, approximately equal, and more than 2-fold higher levels of ONECUT1 protein expression, respectively, compared with Hepa1-6 cells (Figure 1G). In both ONECUT1^int^-EPs and ONECUT1^high^-EPs, the mRNA level of *Alb*, *Afp* and *Ttr*, as well as the secretion of ALB, were significantly reduced compared with those in hepaEP1 cells, suggesting that ONECUT1 expression suppresses hepatic differentiation (Figure 1H, S1D). As *Onecut1* plays a role in biliary development, we further investigated the mRNA expression of cholangiocyte markers. *Krt19* showed exclusive upregulation in ONECUT1^high^-EPs (Figure 1I).

Unexpectedly, downregulation of hepatic markers and upregulation of cholangiocyte markers were also observed in Hepa1-6 cells transfected with *Onecut1* shRNA (Figure S2A, S2B). Moreover, KRT19 protein expression was detected in ONECUT1-knockdown cells but not in ONECUT1^high^-EPs with *Krt19* upregulation, indicating impaired cholangiocyte maturation in ONECUT1-overexpressing cells (Figure S2C, S2D, S2E).

In summary, gradual overexpression of ONECUT1 in ONECUT1^low^-EPs causes suppression of hepatic markers and a greater dose-dependent increase in *Krt19* mRNA expression, resulting in ONECUT1^low^*Alb^high^Krt19^low^* hepatocyte-like cells, ONECUT1^int^*Alb^low^Krt19^low^* dedifferentiated cells, and ONECUT1^high^*Alb^low^Krt19^high^* cholangiocyte-like cells, which collectively represent an incomplete multistate hepatocyte-to-cholangiocyte transdifferentiation process.

### Single-cell RNA-seq reveals the LPC transition during hepatocyte-to-cholangiocyte transdifferentiation

Recently, a single-cell RNA-seq study (Merrell et al. 2021) reported more advanced hepatocyte-to-cholangiocyte transdifferentiation in 3,5-diethoxycarbonyl-1,4-dihydrocollidine (DDC)-treated *Smad4* mutant mice. However, cells undergoing transdifferentiation were treated as a single cluster in the original report. To monitor the transcriptome kinetics during hepatocyte-to-cholangiocyte transdifferentiation, we tried to reanalyze the dataset at a higher resolution.

Single-cell RNA-seq data were obtained from GSE157698, in which *Rosa^YFP/YFP^Smad4^fl/+^* control mice and *Rosa^YFP/YFP^Smad4^fl/fl^* mutant mice were injected with hepatocyte-specific AAV8-TBG-Cre and were then treated with DDC. A total of 23 clusters were identified at a resolution of 0.9 with Seurat (Hao et al. 2021) (Figure S3A). As the KRT19 protein was not expressed in ONECUT1^high^*Alb^low^Krt19^high^* cells, mature cholangiocytes were defined as a *Krt19^+^* cluster present in control bulk liver parenchyma but not in control YFP^+^ cells due to incomplete hepatocyte-to-cholangiocyte transdifferentiation (Figure S3B). After definition of cholangiocytes, *eYFP^-^* cells, contaminating hepatic stellate cells (cluster 11 and 21), endothelial cells (cluster 22) and Kupffer cells (cluster 20) were omitted from further analysis (Figure S3C). *eYFP^+^* hepatocyte-derived cells, including the estimated mature cholangiocytes (cluster 6), formed a continuum rather than discrete clusters (Figure 2A).

**Figure 2.**
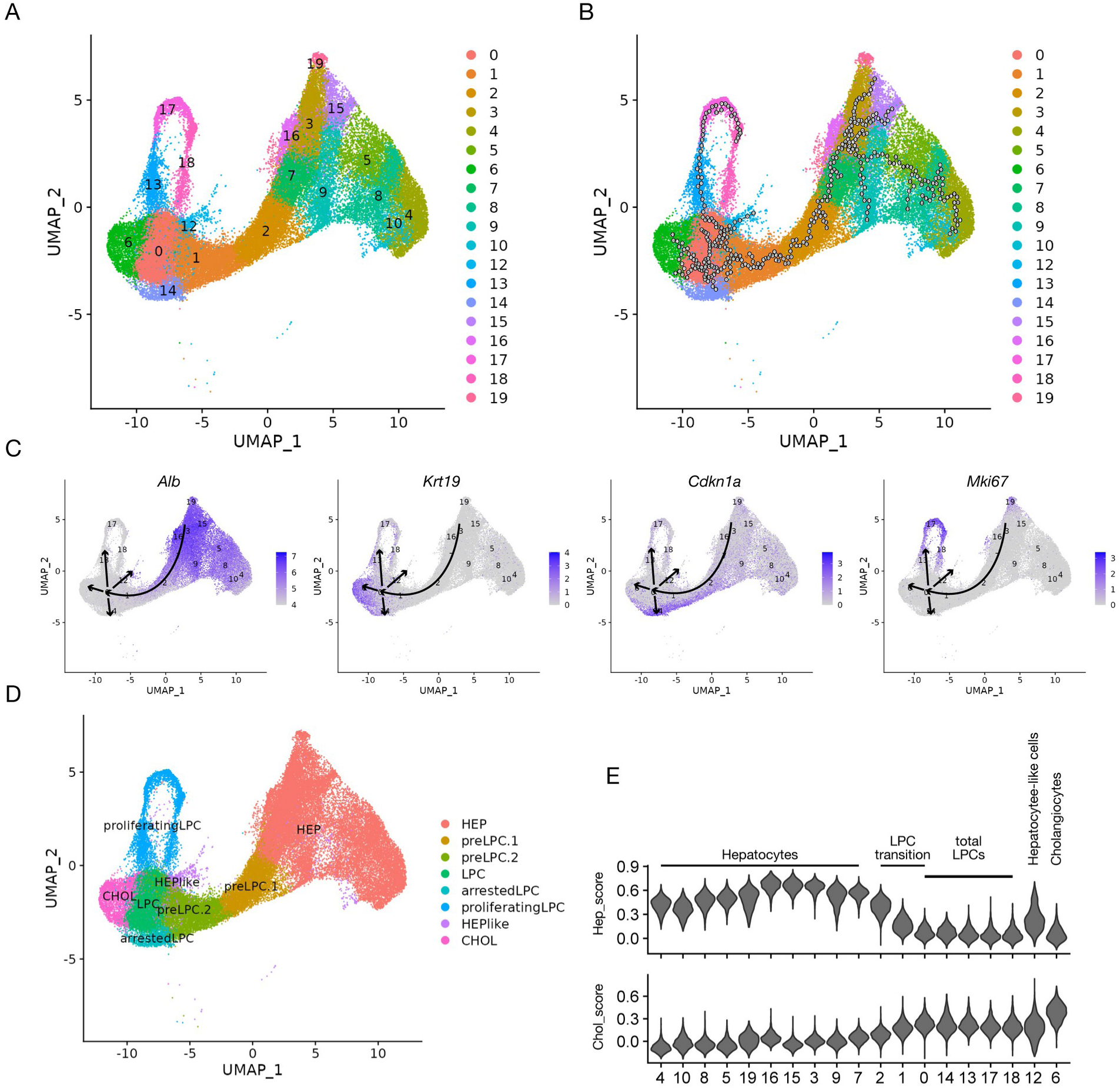
Hepatocyte-to-cholangiocyte transdifferentiation continuum. **(A)** Single-cell RNA-seq profiling of hepatocyte-derived cells by cluster. **(B)** Trajectory analysis of hepatocyte-derived cells. **(C)** Expression of *Alb*, *Krt19*, *Cdkn1a* and *Mki67* in hepatocyte-derived cells. **(D)** Hepatocyte-to-cholangiocyte transdifferentiation continuum by cell type. **(E)** Relative scores of hepatic and cholangiocyte markers by cluster.

Trajectory analysis with Monocle (Qiu et al. 2017) and differentially expressed gene (DEG) analysis suggested that hepatocytes transit along clusters 2, 1, and 0 and then branch into *Alb^high^* hepatocyte-like cells (cluster 12), *Krt19^+^* estimated mature cholangiocytes (cluster 6), *Cdkn1a^+^* arrested cells (cluster 14), and proliferating cells (clusters 13, 17, and 18), indicating the bipotent-liver-progenitor-cell-like ability and self-renewal capacity of the cells in cluster 0 (Figure 2B, 2C). According to the results, hepatocyte-to-cholangiocyte transdifferentiation comprises a continuum of 3 major states: hepatocytes, transitioning LPCs (including preLPC.1s, preLPC.2s, and total LPCs, which are further composed of LPCs, arrested LPCs and proliferating LPCs), and terminally differentiated cells (hepatocyte-like cells or cholangiocytes) (Figure 2D). To assess the transdifferentiation level, a hepatic score (hep_score) and a cholangiocytic score (chol_score) were determined by analysis of hepatic and cholangiocyte markers, respectively (Franzén, Gan, and Björkegren 2019) (Figure S4). Along the transdifferentiation continuum, the chol_score increased during the LPC transition, while the hep_score transiently increased before the LPC transition and then immediately decreased during the LPC transition. Transdifferentiated hepatocyte-like cells showed higher hep_score values than LPCs (Figure 2E).

To confirm that *Smad4* mutation affects advanced hepatocyte-to-cholangiocyte transdifferentiation, we compared the percentages of the continuum components between control mice and *Smad4* mutant mice. Consistent with the original report, accumulation of total LPCs and transdifferentiated cholangiocytes was observed in *Smad4* mutant mice (Figure S5A, S5B). To assess the enhancement of transdifferentiation, we further calculated transition rates as the ratios of arrested LPCs, proliferating LPCs, hepatocyte-like cells or cholangiocytes to LPCs. Compared with control mice, *Smad4* mutant mice showed a decreased LPC-hepatocyte-like cell transition rate and a similar LPC-cholangiocyte transition rate. On the other hand, the transition of arrested LPCs was suppressed and the transition of proliferating LPCs was enhanced in *Smad4* mutant mice, indicating promoted proliferation of LPCs (Figure S5C). These results demonstrate that the accumulation of cholangiocytes in *Smad4* mutant mice more likely resulted from an increase in the LPC number due to enhanced self-renewal than due to advanced transdifferentiation.

### *Onecut1* accumulates throughout the LPC transition

As *Smad4* is considered to mainly affect proliferation of LPCs, only data from control mice were used in further analyses. In control mice, *Onecut1^+^* cells were observed in only a small proportion of hepatocytes, with expansion before and throughout the LPC transition. The transdifferentiated hepatocyte-like cell component showed almost complete loss of *Onecut1^+^* cells, while the cholangiocyte component showed mild loss of *Onecut1^+^* cells (Figure 3A). These results suggest a potential role of *Onecut1* during the LPC transition. In *Onecut1^+^* cells extracted from enriched clusters, *Onecut1* was gradually upregulated from hepatocytes to LPCs and was then downregulated in cholangiocytes, accompanied by *Alb* downregulation and *Krt19* upregulation, showing a pattern similar to that in ONECUT1-EPs (Figure 3B). Compared to those in *Onecut1^-^* cells, preLPC.1 component, preLPC.2 component and LPC component were increased in *Onecut1^+^* cells, indicating that *Onecut1* facilitates the LPC transition (Figure 3C, 3D). These results are consistent with the gradual overexpression of ONECUT1 in ONECUT1^low^-EPs causing incomplete hepatocyte-to-cholangiocyte transdifferentiation. However, no marked differences in the levels of hepatic or cholangiocyte markers were observed between *Onecut1^-^* and *Onecut1^+^* cells of the same cell type (Figure 3E).

**Figure 3.**
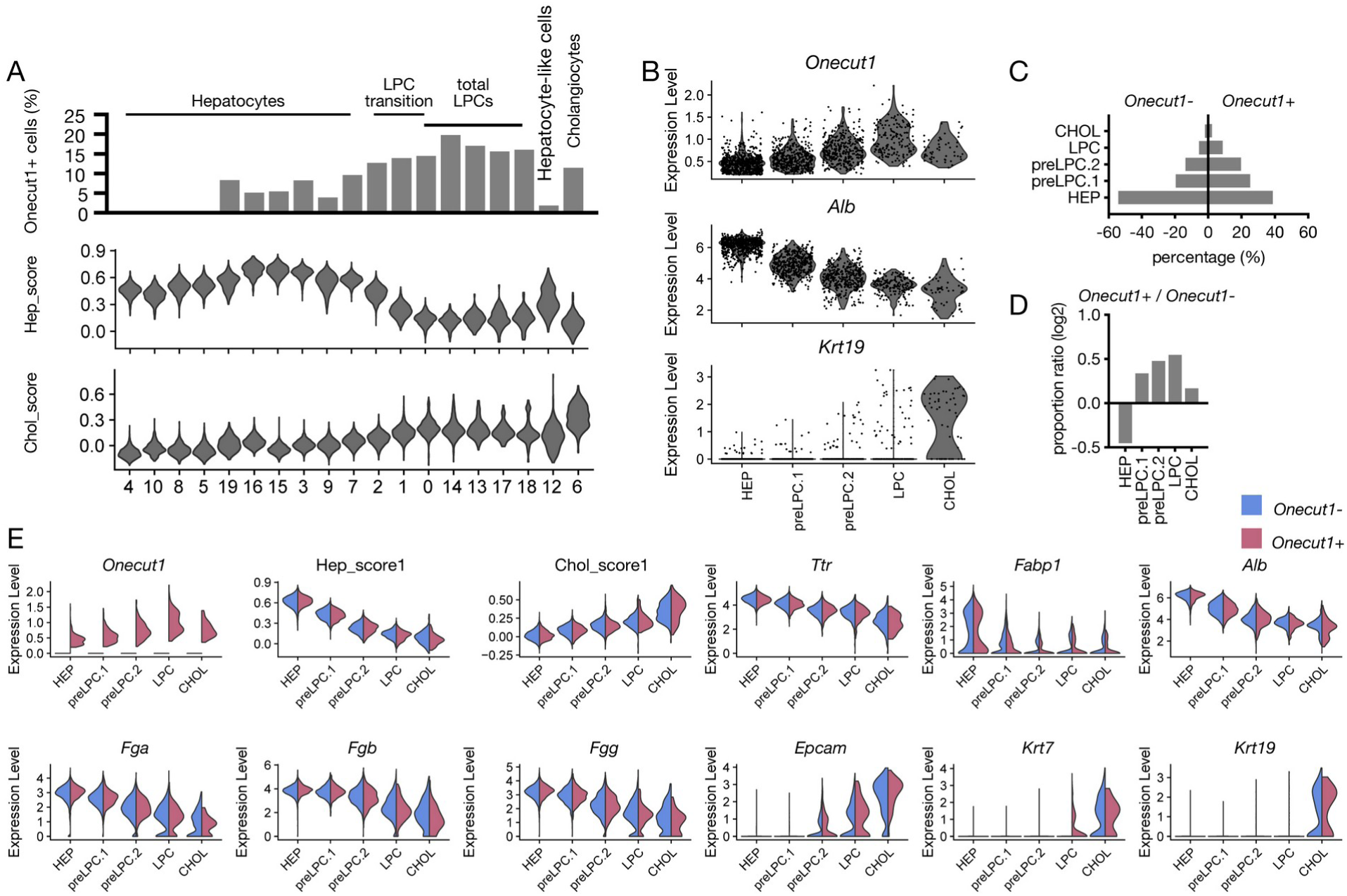
Enrichment and expression of *Onecut1* during the LPC transition. **(A)** Accumulation of *Onecut1^+^* cells with hep_score and chol_score values in control mice by cluster. **(B)** Expression of *Onecut1* in *Onecut1^+^* cells extracted from enriched clusters by cell type. **(C)** Proportions of *Onecut1^-^* and *Onecut1^+^* cells by cell type. **(D)** Ratios of proportions of components in *Onecut1^+^* cells to those in *Onecut1^-^* cells by cell type. **(E)** Expression of the indicated genes in *Onecut1^-^* and *Onecut1^+^* cells by cell type.

### ONECUT1^int^*Alb^low^Krt19^low^* dedifferentiated cells show high pulmonary metastatic potential

As LPCs are considered the origin of metastasis-initiating cells, we performed an *in vivo* metastasis assay to examine the metastatic potential of ONECUT1-EPs. Twenty-four days after intravenous injection, pulmonary metastases derived from ONECUT1^int^-EPs and Hepa1-6 cells but not from ONECUT1^low^-EPs or ONECUT1^high^-EPs were observed (Figure 4A, 4B). To confirm that high overexpression of ONECUT1 inhibits metastasis, we further introduced 3FLAG-ONECUT1 into metastatic Hepa1-6 cells or ONECUT1-EP2s (Figure 4C). Both generated cell lines showed downregulation of *Afp* mRNA expression, upregulation of *Krt19* mRNA expression, and decreased metastatic potential compared to the corresponding parental cells (Figure 4D, 4E, 4F, 4G). These results indicate that only intermediate ONECUT1-expressing cells with dedifferentiated traits acquire metastatic potential.

**Figure 4.**
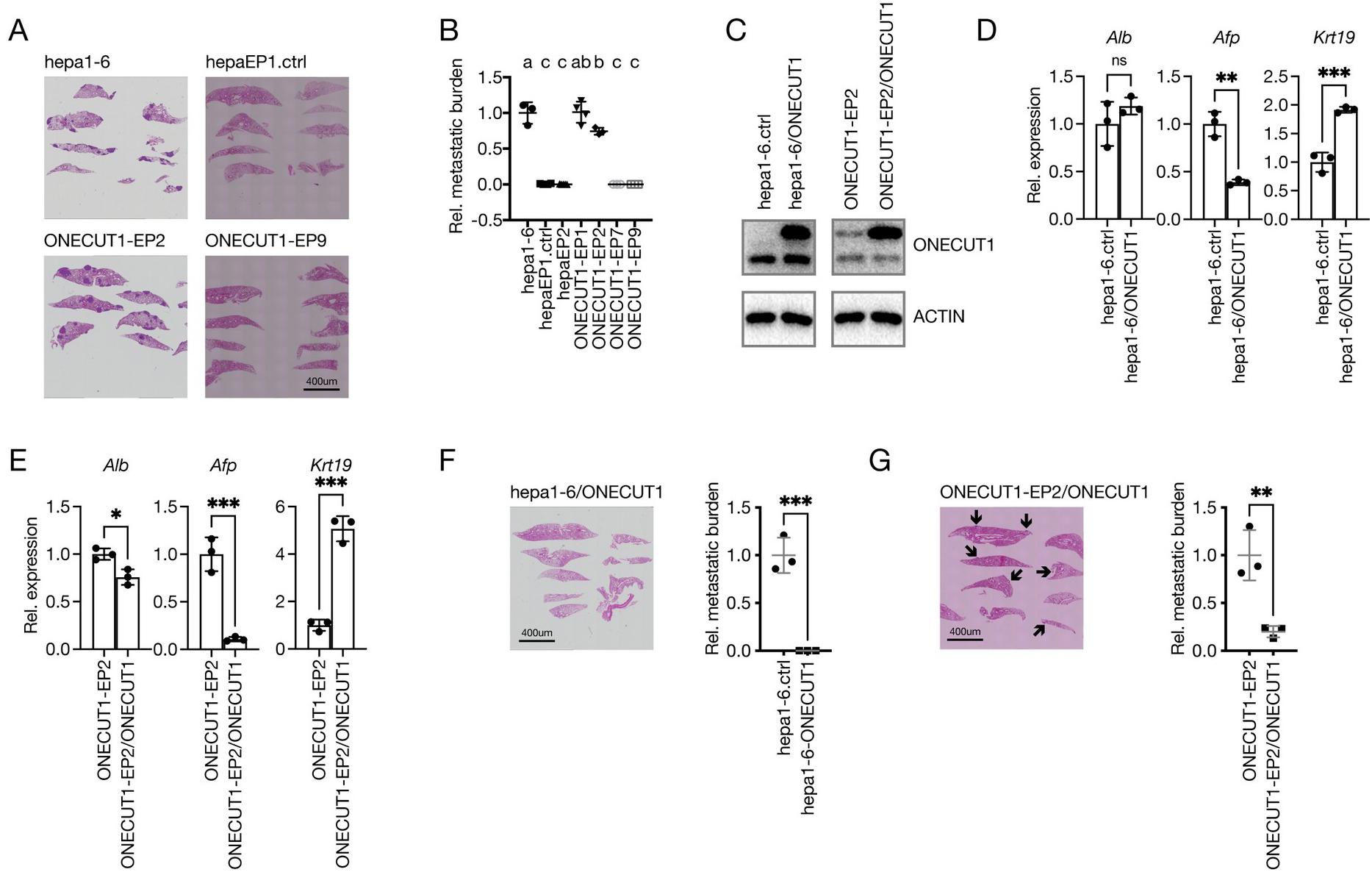
ONECUT1^int^*Alb^low^Krt19^low^* cells show high pulmonary metastatic potential. **(A)** Hematoxylin and eosin staining of lungs 24 days after intravenous injection. **(B)** Relative lung metastatic burden. The hepa1-6 metastatic burden was set to 1; n = 3 for hepa1-6, ONECUT1-EP2, ONECUT1-EP7 and ONECUT1-EP9; n = 4 for hepaEP1 and ONECUT1-EP1; n = 5 for hepaEP2; the different letters indicate significant differences by 1-way ANOVA. **(C)** Immunoblotting of ONECUT1 (upper) and ACTIN (lower). (D) Expression of *Alb*, *Afp* and *Krt19*, as determined by quantitative RT‒ PCR. The mRNA level of Hepa1-6 was set to 1; n = 3; * indicates a significant difference by a t test. **(E)** Expression of *Alb*, *Afp* and *Krt19*, as determined by quantitative RT‒ PCR. The ONECUT1-EP2 mRNA level was set to 1; n = 3; * indicates a significant difference by a t test. **(F)** Hematoxylin and eosin staining of lungs and calculation of the relative metastatic burden. The Hepa1-6 metastatic burden was set to 1; n = 3; * indicates a significant difference by a t test. **(G)** Hematoxylin and eosin staining of lungs and calculation of the relative metastatic burden. The ONECUT1-EP2 metastatic burden was set to 1; n = 3; * indicates a significant difference by a t test.

### LPCs exhibit a “markerless” phenotype

None of the reported LPC or stem cell markers distinguished preLPC.1s, preLPC.2s or LPCs along the hepatocyte-to-cholangiocyte transdifferentiation continuum (Figure S6A). To identify the possible origin of metastasis-initiating cells, we extracted markers from single-cell RNA-seq data. However, even with relaxed cutoff criteria (log2FC > 0.5 & p < 0.05), no preLPC.2 markers were extracted. *Mup20*, *Hsd17b13*, *Mug1*, *Csad*, *Itih3*, *Slc2a2* and *Hpx* were extracted as preLPC.1 markers, indicating that these genes reflect more hepatocyte-like traits. Only *Vim* was extracted as an LPC marker. Although *Csad* and *Vim* showed lower background expression in hepatocyte-derived cells, their expression was observed in neighboring clusters (Figure S6B, S6C).

As *Vim* is a key marker of epithelial-mesenchymal transition (EMT), we further assessed the expression of EMT marker genes. An epithelial score (epi_score) and a mesenchymal score (mes_score) were determined by analysis of epithelial and mesenchymal markers, respectively (Tan et al. 2014). Along the transdifferentiation continuum, the epi_score gradually increased, and the mes_score increased during the LPC transition and remained increased in transdifferentiated cells. Compared to those of LPCs, the epi_score values of transdifferentiated hepatocyte-like cells were similar, while those of cholangiocytes were increased, suggesting that cholangiocyte maturation is associated with possible mesenchymal-epithelial transition (MET). No decrease in the epi_score or loss of *Cdh1* was observed during hepatocyte-to-cholangiocyte transdifferentiation (Figure S6D). Among the core EMT transcription factors, only *Zeb2* was upregulated in a small proportion of LPCs (Figure S6E).

It is reasonable that no specific markers were extracted, as the continuum indicates seamless transition states rather than different cell types. We further monitored the continuous nature of gene expression during the LPC transition by trajectory analysis. Along pseudotime, preLPC.1s, preLPC.2s, LPCs and cholangiocytes overlapped with the neighboring clusters (Figure S7A). The expression levels of *Cd24a* and *Spp1* showed early increases, and those of *Vim*, *Cd44*, *Epcam*, *Sox4* and *Sox9* increased later. At the end of the continuum, *Sox9* and *Spp1* expression decreased, while *Krt19*, *Krt7* and *Tacstd2* (also known as *Trop2*) expression rapidly increased (Figure S7B).

### Hepatically differentiated tumor cells acquire metastatic potential in the early stage of the LPC transition

To investigate whether distant metastases show preLPC.2-like or LPC-like traits, we analyzed gene expression in distant metastases of human HCC. Microarray data were obtained from GSE40367 (Roessler et al. 2015). Normal liver samples were used as controls due to the complex heterogeneity of the primary tumors. A nonsignificant increase in *KRT19* expression was found in the distant metastases (Figure 5A). As the distant metastases were clearly separated into two discrete groups with low and high *KRT19* expression, we further investigated the difference among normal liver tissues, *KRT19^low^* distant metastases and *KRT19^high^* distant metastases. Compared to those in normal liver tissues, the expression levels of *ALB* and the preLPC.1 marker *CSAD* were decreased in both *KRT19^low^* and *KRT19^high^* metastases, while the expression levels of *KRT19* and the LPC marker *VIM* were increased exclusively in *KRT19^high^* metastases (Figure 5B). Seven genes were further extracted with significant differences between *KRT19^low^* and *KRT19^high^* metastases and between preLPC.2s and LPCs. The expression of all genes increased along transdifferentiation pseudotime and also upregulated in *KRT19^high^* metastases. Notably, *Rgs5* reached peak expression in the LPC state, indicating the presence of preLPC.2-like traits in KRT19^low^ metastases (Figure 5C). The results revealed the existence of both *ALB^low^KRT19^low^VIM^low^* and *ALB^low^KRT19^high^VIM^high^* distant metastases, suggesting that metastatic potential is acquired during the LPC transition in a state before upregulation of *KRT19* and *VIM*.

**Figure 5.**
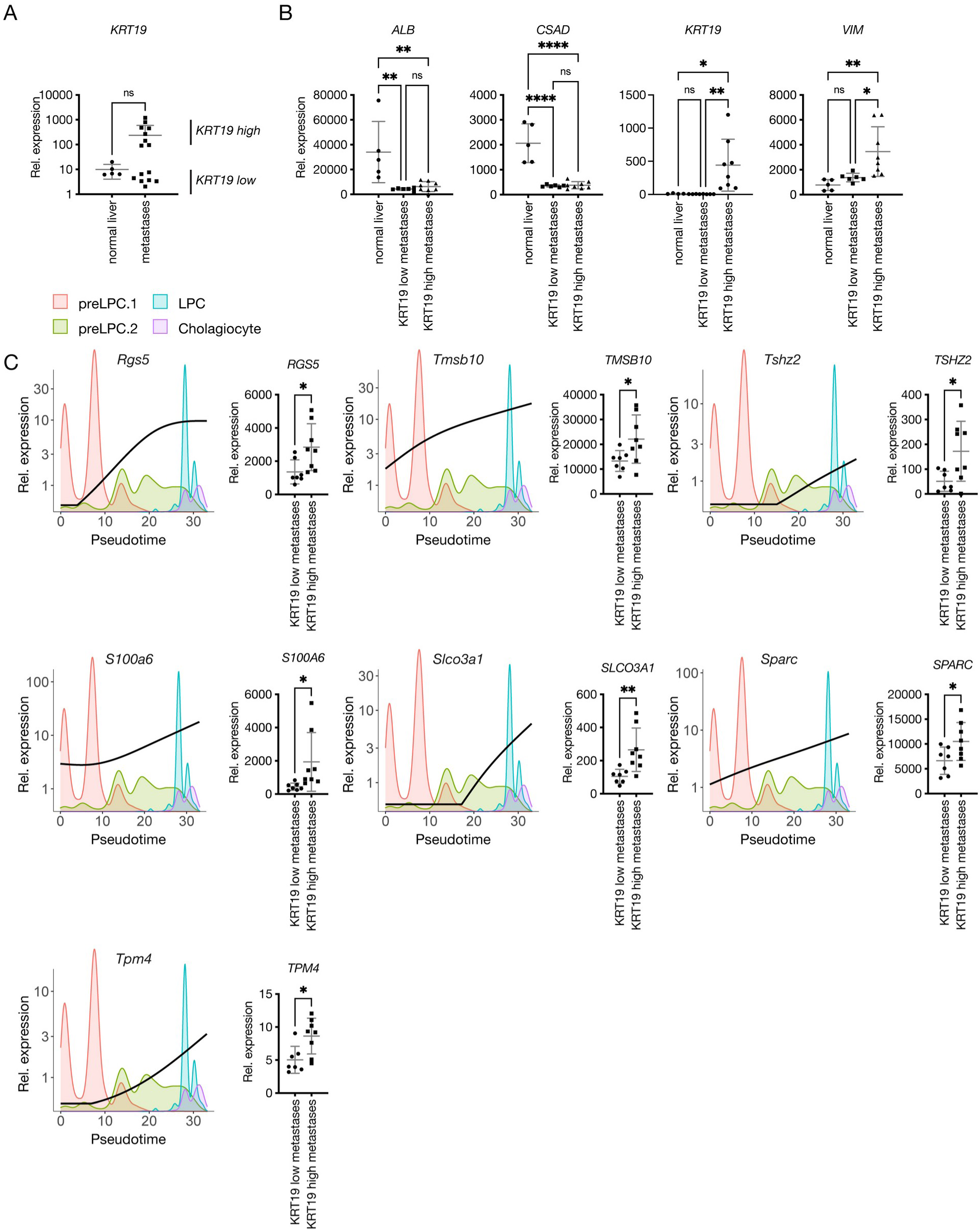
PreLPC.2-like and LPC-like distant metastases in human HCC. **(A)** Expression of *KRT19* in normal liver tissue and distant metastases. Nonsignificant, t test. **(B)** Expression of *ALB*, *CSAD*, *KRT19* and *VIM* in normal liver tissue, *KRT19^low^* metastases and *KRT19^high^* metastases. Outliers were removed using Tukey’s method; * indicates a significant difference by 1-way ANOVA. **(C)** Expression of common DEGs shown as an expression curve along pseudotime (left) or by expression in human HCC data (right). * indicates a significant difference by a t test.

Main analyses were also performed on only sorted YFP^+^ cells, and similar results were observed (Figure S8-10).

## Discussion

We tried to identify LPCs with single-cell RNA-seq data of eYFP-labeled hepatocyte-derived cells from DDC-treated control and *Smad4* mutant mice. The hepatocyte-derived cells formed a continuum from *Alb^high^* cells to *Alb^low^Krt19^+^* cells, indicating the seamless trajectory of hepatocyte-to-cholangiocyte transdifferentiation. Trajectory analysis revealed that a single cluster branched into *Alb^high^* hepatocyte-like cells, *Krt19^+^* cholangiocytes and proliferating cells, suggesting LPC-like traits. We demonstrated that hepatocytes transition into LPCs and then further differentiate into hepatocyte-like cells or cholangiocytes. Consistent with the report that SPP1^+^MKI67^+^ hepatocyte-derived reactive ductules were KRT19^-^, proliferating cells diverged from LPCs but not cholangiocytes. The proportion of proliferating LPCs was markedly increased in *Smad4* mutant mice, while the ratio of cholangiocytes to LPCs was similar in control and *Smad4* mutant mice, indicating that the accumulation of cholangiocytes in *Smad4* mutant mice resulted from enhanced self-renewal rather than differentiation of LPCs.

Consistent with reports from other groups, the common LPC markers failed to distinguish LPCs. In the single-cell RNA-seq data, only *Vim* was extracted as an LPC marker, and *Vim* was also expressed in neighboring clusters and a wide range of cell types, indicating the difficulty of identifying LPCs with markers *in vivo*. Although expression of *Vim* suggests the presence of transient mesenchymal traits during hepatocyte-to-cholangiocyte transdifferentiation, no typical EMT was observed.

By gradually expressing ONECUT1 in Hepa1-6-derived ONECUT1^low^*Alb^high^Krt19^low^* hepatocyte-like cells, we obtained ONECUT1^int^*Alb^low^Krt19^low^* dedifferentiated cells and ONECUT1^high^*Alb^low^Krt19^high^* cholangiocyte-like cells with downregulation of *Alb* and upregulation of *Krt19*. Although *Krt19* mRNA expression was upregulated in ONECUT1^high^*Alb^low^Krt19^high^* cells, KRT19 protein expression was not detected, indicating impaired cholangiocyte maturation. On the other hand, knockdown of *Onecut1* in Hepa1-6 cells facilitated KRT19 protein expression, consistent with the premature cholangiocyte differentiation observed in *Onecut1* knockout mice. During development, ONECUT1 is weakly expressed in the liver parenchyma and accumulates in the ductal plate, and ONECUT1 expression is then completely lost in mature cholangiocytes. Single-cell RNA-seq analysis revealed that both the accumulation of *Onecut1^+^* cells and the level of *Onecut1* expression in *Onecut1^+^* cells increased before and throughout the LPC transition and then decreased in cholangiocytes. As similar behaviors were observed in all HCC cell models, during development and in DDC-treated mice, we demonstrated that *Onecut1* expression facilitates the LPC transition, which cannot be simply reversed by knockdown *Onecut1*. *Onecut1* expression in LPCs needs to be correctly regulated to determine cholangiocyte maturation, probably through TGFb signaling pathways (Clotman et al. 2005). Although ablation of *Smad4* resulted in accumulation of cholangiocytes, we believe that this effect is dependent mainly on disruption of *Smad4*-mediated growth inhibition.

Despite the accumulation of preLPC.1s, preLPC.2s and LPCs among *Onecut1^+^* cells, no marked differences in gene expression were observed between *Onecut1^-^* and *Onecut1^+^* cells of the same cell type. Thus, the generation of *Onecut1^-^* cells could either be regulated by other pathways or result from expansion of *Onecut1^+^* cells with *Onecut1* loss (although *Onecut1* loss is suggested to be associated with cholangiocyte maturation), and these possibilities need further investigation. According to the results, we presume that cells of the same cell type show similar traits regardless of whether *Onecut1* is expressed. Thus, we assume that the ONECUT1-EP model could partially represent the LPC transition and not only *Onecut1^+^* cells *in vivo*.

As LPCs are considered the origin of metastasis-initiating cells, we investigated the metastatic potential of ONECUT1-EPs and found that only ONECUT1^int^*Alb^low^Krt19^low^* dedifferentiated cells formed distant metastases. In human HCC microarray data, distant metastases showed both *ALB^low^KRT19^low^VIM^low^* and *ALB^low^KRT19^high^VIM^high^* phenotypes. These results revealed that hepatically differentiated tumor cells acquire metastatic potential before *KRT19* and *VIM* upregulation, probably in the preLPC.2 state. Although not suitable for identification, the expression of cholangiocyte or mesenchymal markers in hepatocyte-derived cells could still be used to present the existence of preLPC.2 cells with metastatic potential. On the other hand, the metastatic potential of LPCs has not been excluded due to the formation of *ALB^low^KRT19^high^VIM^high^* metastases, which could result from either LPC-derived metastases or further LPC transition of preLPC.2 or preLPC.2-derived metastases (Figure S11).

The LPC transition formed a seamless continuum, with most DEGs expressed in the intermediate state between hepatocytes and cholangiocytes. Our study emphasizes the importance of *in vitro* studies due to the markerless phenotype of the preLPC.2s, which are considered candidate metastasis-initiating cells. However, the multistate ONECUT1-EP model could not correctly reproduce the gradual transition and showed high expression variability among the cell lines. We also tried to generate inducible cell lines but failed possibly due to the leakiness of ONECUT1. To further investigate metastatic characteristics during the LPC transition and determinants of terminal differentiation, sensitively controlled inducible cell models need to be established.

## Notes

### Competing Interest Statement

The authors have declared no competing interest.

### Summary of Updates

The parameters of the single-cell RNA-seq analyses were improved.

