## supplementary_figure for "*Onecut1* partially contributes to the liver progenitor cell transition and acquisition of metastatic potential in hepatocellular carcinoma"

### Materials and Methods

**Cell lines.** All cell lines were maintained in DMEM containing 10% FBS at 37 °C in 5% CO2. Cells were reseeded at 3 x $10^4$ cells/cm2 every 48 h. Stable clones were established by transfection of cells with pIRESEF1α/ONECUT1, pIRESEF1α/3FLAG-ONECUT1, pSuper+/shONECUT1 or corresponding empty vectors using jetPEI (Polyplus, FR) and selection with neomycin (100-800 μg/ml), puromycin (0.5-4 μg/ml) or hygromycin (200-1600 μg/ml).

**Immunoblotting and immunoprecipitation.** For immunoblotting of cell lysates, cells were lysed in RIPA buffer. For immunoblotting of culture supernatants, the regular culture medium was replaced with serum-free DMEM. Serum-free supernatants were collected 16 h later and concentrated with Amicon Ultra filters (MWCO, 3k, Merck Millipore, US). For immunoprecipitation, culture supernatants were collected after 48 h of incubation and filtered through 0.22-μm filters. Immunoprecipitation was performed with an anti-ALB antibody (A90-134F-3, Bethyl, US) and protein G Sepharose (GE Healthcare, US). Antibodies against ALB (ab207327, Abcam, UK), ONECUT1 (ab186743, Abcam, UK), KRT19 (ab52625, Abcam, UK) and ACTIN (MAB1501, Merck Millipore, US) were used for immunoblotting. CBB Stain One (Nacalai Tesque, JP) was used for total protein staining.

**RNA isolation and quantitative reverse transcription (RT)–PCR.** Total RNA was extracted with Sepasol-RNA I Super G (Nacalai Tesque, JP) prior to reverse transcription using ReverTra Ace qPCR RT Master Mix (Toyobo, JP) and purification by ethanol precipitation. mRNA expression was measured by real-time PCR using

THUNDERBIRD SYBR qPCR Mix (Toyobo, JP) and a StepOnePlus Real-Time PCR System (Applied Biosystems, US) and was then normalized to GAPDH expression.

**Immunofluorescence analysis.** For immunofluorescence analysis, cells were cultured in 24-well dishes for 48 or 72 h and fixed with 10% formalin prior to permeabilization with 0.5% Triton X-100. After blocking in 5% BSA, the cells were incubated sequentially with an anti-KRT19 antibody (ab52625, Abcam, UK) overnight at 4 °C, TRITC-labeled anti-rabbit IgG for 1 h at room temperature, and a FITC-labeled anti-ALB antibody (A90-134F-3, Bethyl, US) overnight at 4 °C. All images were acquired using a BZ-9000 BIOREVO fluorescence microscope (Keyence, JP).

**Mice, xenograft assay and *in vivo* metastasis assay.** BALB/c nude mice (male, 6 weeks old; SLC, JP) were used for in vivo studies. For the xenograft assay, 2 x $10^6$ cells were injected into the flanks of nude mice. Tumor volume was calculated as Length x $(\text{Width})^2$ / 2. For the *in vivo* metastasis assay, 2 x $10^6$ cells were injected into the tail veins of nude mice. Lungs were collected after 24 days and were then fixed with 10% formalin. For each mouse, 2 or 3 sections sliced at 0.5 mm intervals were used for hematoxylin and eosin staining. Metastatic burden was calculated as average of (counts of metastatic foci)/(tissue area). All animal experiments were reviewed and approved by the Animal Committee of Kyoto University.

**Single-cell RNA-seq analysis.** Filtered gene data were obtained from GSE157698. Low-quality cells and genes were filtered with the following criteria: min.cells = 3, 200 < nFeatures < 6000 and percent.MT < 30. Data for bulk liver parenchyma from control mice (GSM4885630 and GSM4885634), *eYFP*$^+$ bulk liver parenchyma from Smad4 mutant mice (GSM4885631 and GSM4885635, eYFP > 0), and sorted YFP$^+$ cells from both control mice (GSM4885632 and GSM4885636, *eYFP* > 0) and *Smad4* mutant mice (GSM4885633 and GSM4885637, *eYFP* > 0) were integrated with Seurat. RunUMAP visualization was used for clustering at a resolution of 0.9. Monocle3 and Monocle2 were used for trajectory analysis of the whole continuum and the LPC transition, respectively.

**Microarray analysis.** CEL files were obtained from GSE4036723 and processed with Affymetrix Expression Console Software using the MicroArray Suite 5 (MAS5) method. Outliers were removed with Tukey's method.

**Statistics.** Statistical analyses were performed using GraphPad Prism, Microsoft Excel or R. Data are presented as the mean ± s.d. values. Unless otherwise indicated, statistically significant P values are presented as *, $P < 0.05$; **, $P < 0.01$; ***, $P < 0.001$; or ****, $P < 0.0001$.

Figure S1

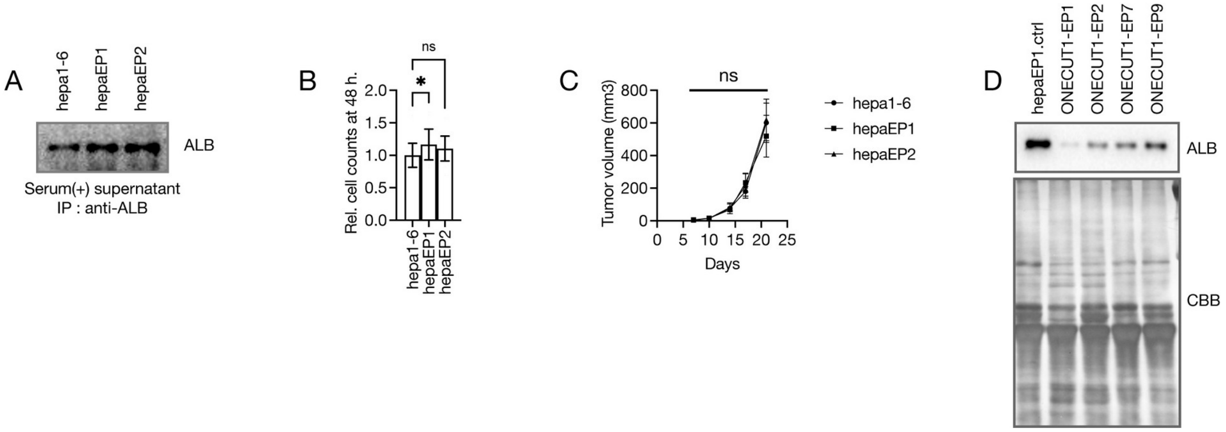


**Figure S1. (A)** Immunoprecipitation of secreted ALB in serum-containing culture supernatants. **(B)** Relative cell counts after 48-h incubation. The count of Hepa1-6 was set to 1; n = 19; * indicates a significant difference by 1-way ANOVA. **(C)** Xenograft assay in nude mice. n = 4; non-significant by 2-way repeated measures ANOVA. **(D)** Immunoblotting of secreted ALB (upper) and total protein (lower).

Figure S2

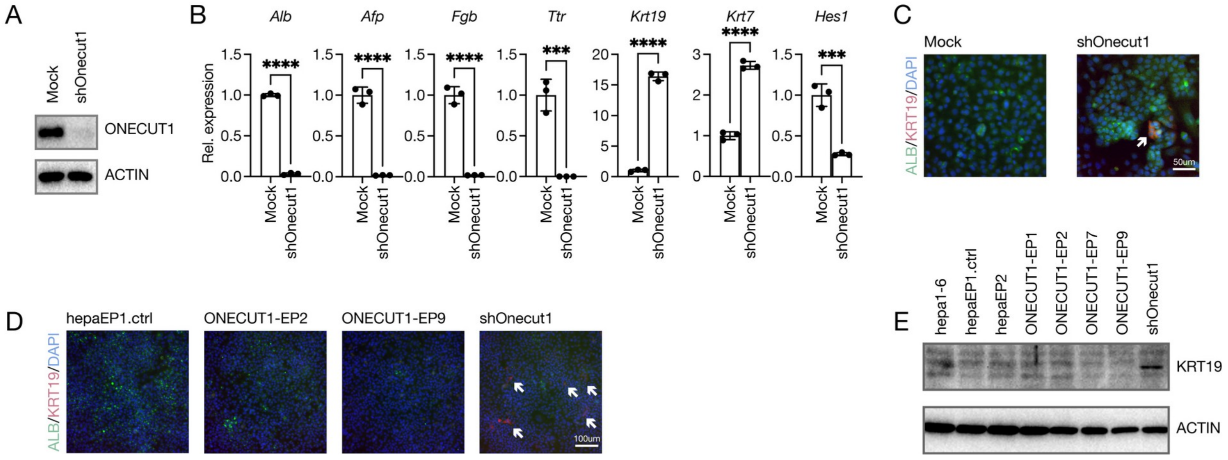


**Figure S2. (A)** Immunoblotting of ONECUT1 (upper) and ACTIN (lower). **(B)** Expression of hepatic and cholangiocyte markers, as measured by quantitative RT–PCR. The mRNA level of Hepa1-6 was set to 1; n = 3; * indicates a significant difference by a t test. **(C, D)** Immunofluorescence staining of ALB (green) and KRT19 (red). **(E)** Immunoblotting of KRT19 (upper) and ACTIN (lower).

Figure S3

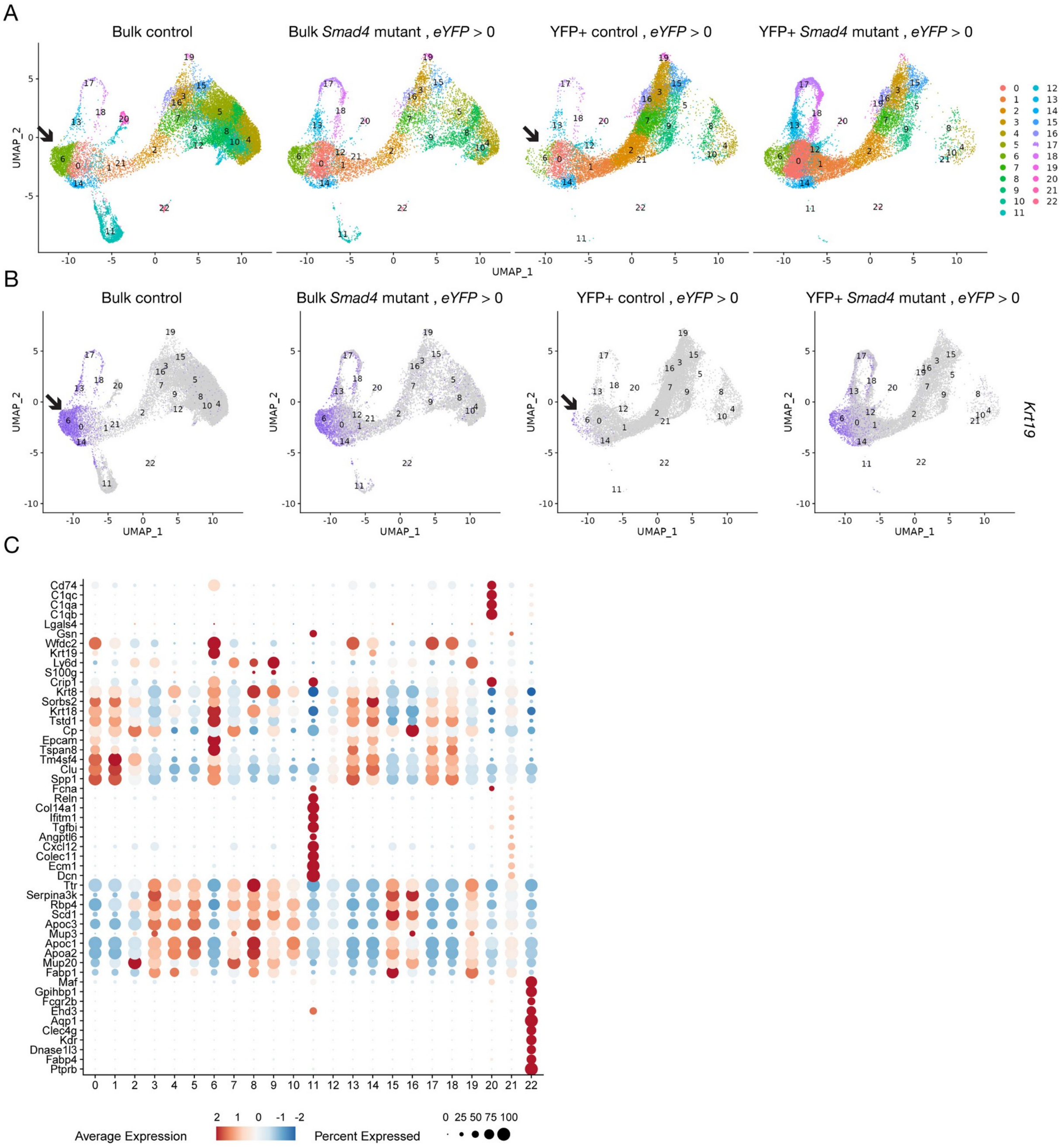


**Figure S3. (A)** Single-cell RNA-seq profiling of hepatocyte-derived cells by cluster. **(B)** Expression of *Krt19* in hepatocyte-derived cells by cluster; the arrows indicate estimated cholangiocyte clusters. **(C)** Expression of hepatic markers, cholangiocyte markers, hepatic stellate cell markers, endothelial cell markers and Kupffer cell markers.

Figure S4

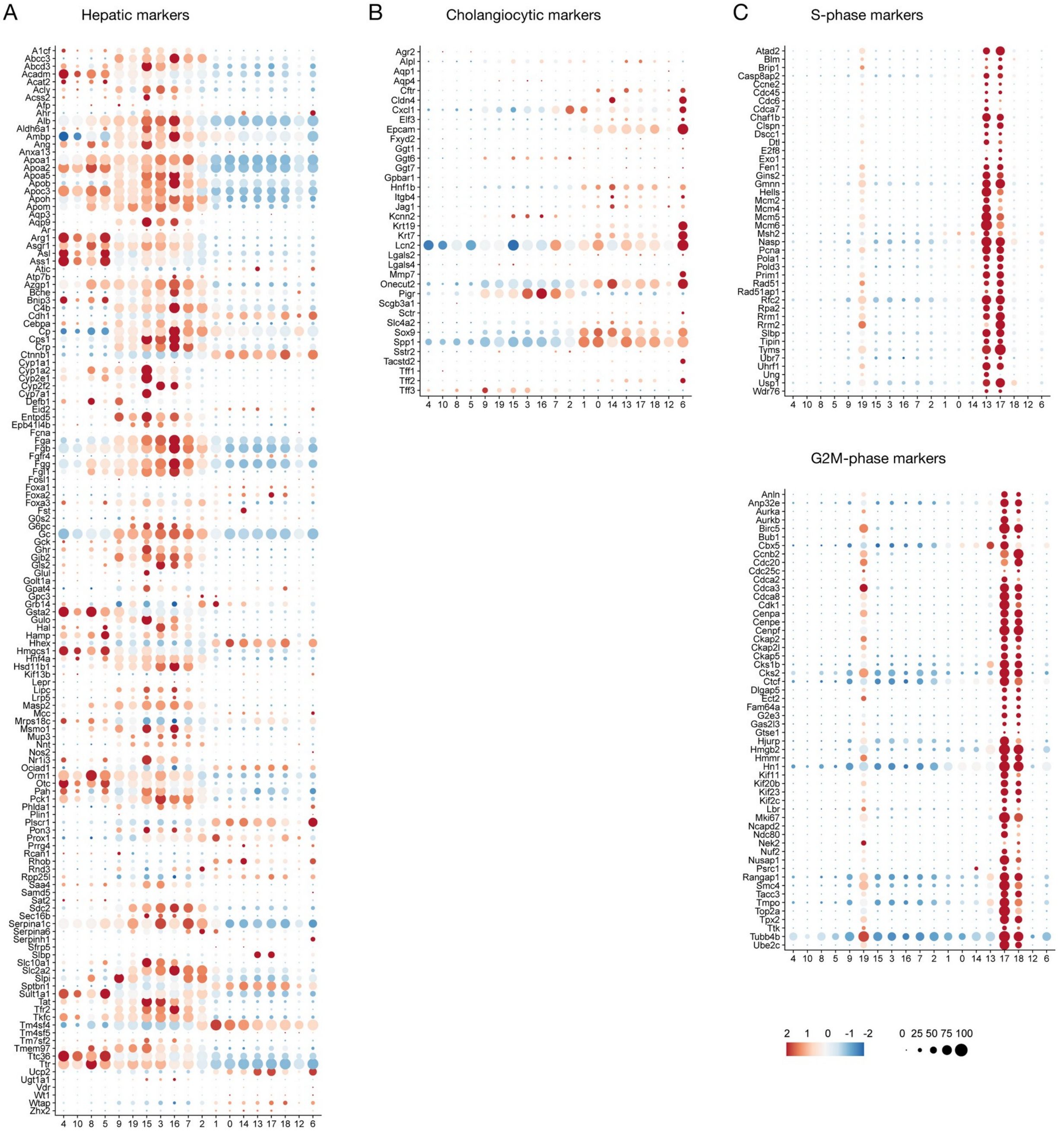


**Figure S4.** Expression of hepatic and cholangiocyte markers used in calculating scores and expression of cell cycle markers.

Figure S5

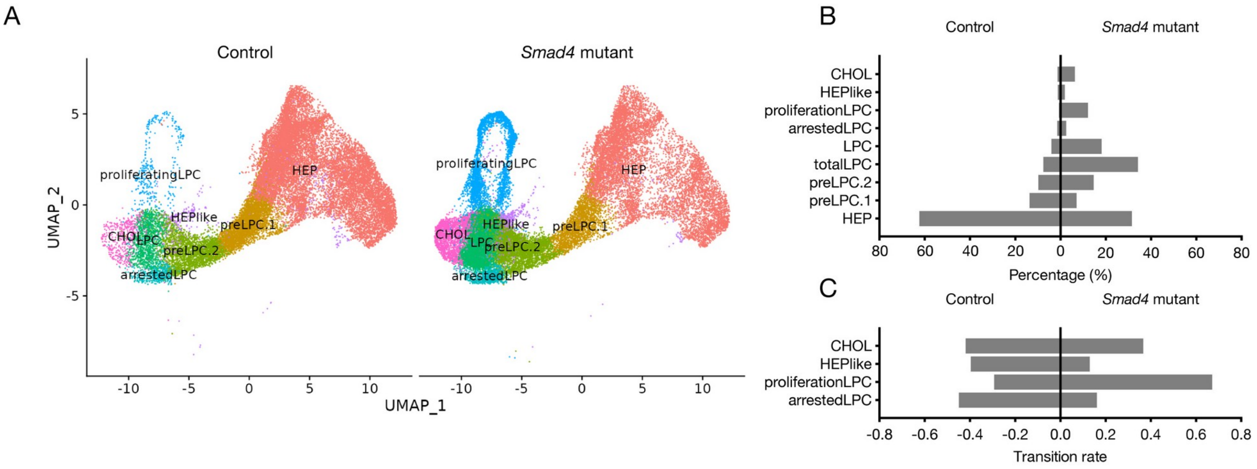


**Figure S5. (A)** Hepatocyte-to-cholangiocyte transdifferentiation continuum in control and *Smad4* mutant mice by cell type. **(B)** Proportions of cells in control and *Smad4* mutant mice by cell type. **(C)** Transition rates in control and Smad4 mutant mice by cell type.

Figure S6

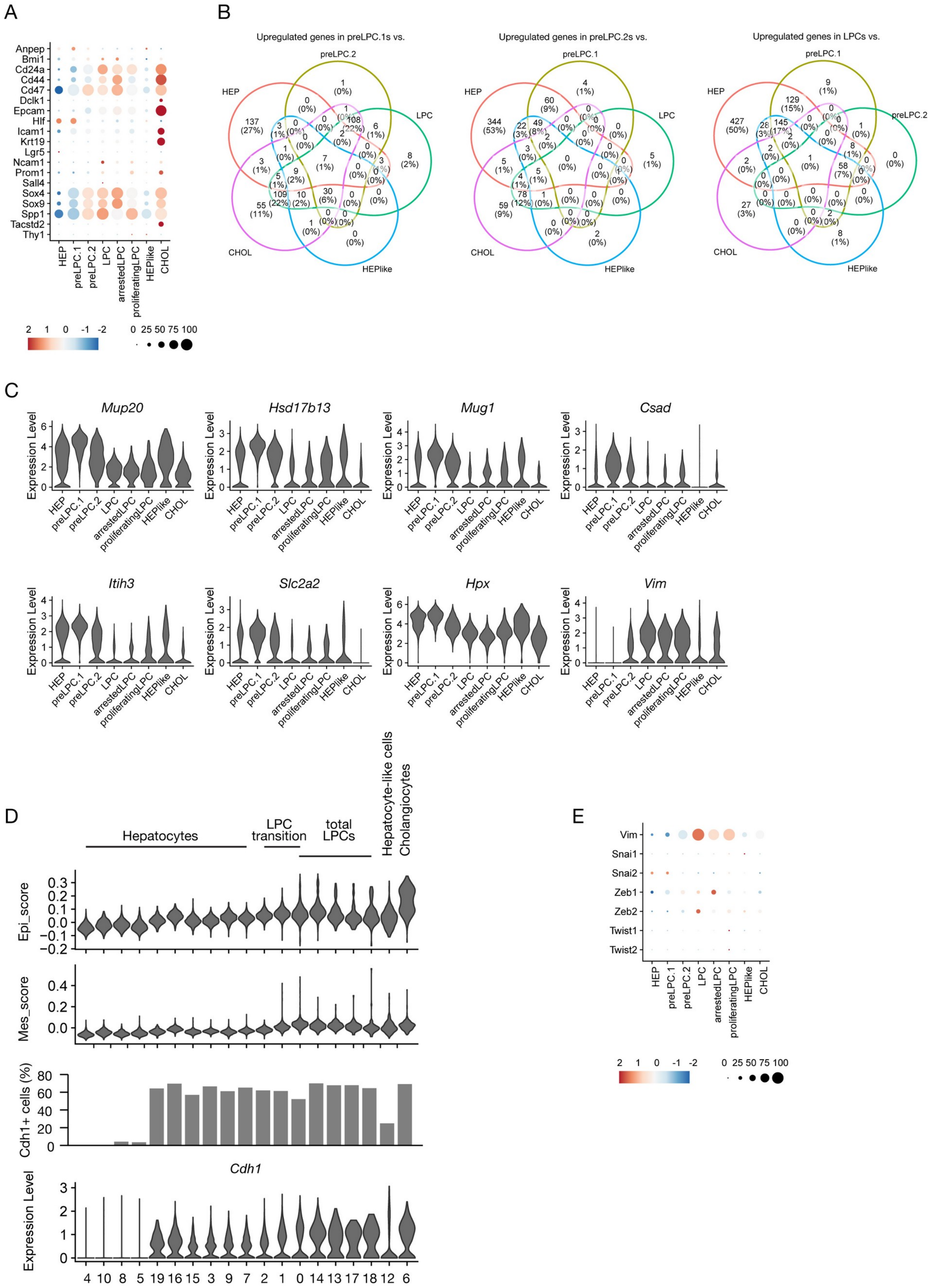

A
B
Upregulated genes in preLPC.1s vs.
Upregulated genes in preLPC.2s vs.
Upregulated genes in LPCs vs.
C
Mup20
Hsd17b13
Mug1
Csad
Itih3
Slc2a2
Hpx
Vim
D
Hepatocytes
LPC transition
total LPCs
Hepatocyte-like cells
Cholangiocytes
Epi_score
Mes_score
Cdh1+ cells (%)
Cdh1
E

**Figure S6. (A)** Expression of reported LPC markers by cell type. **(B)** Venn diagram of upregulated genes in preLPC.1s, preLPC.2s and LPCs. **(C)** Expression of common upregulated genes extracted in (B) by cell type. **(D)** Relative scores of epithelial and mesenchymal markers and expression of *Cdh1* and *Cdh2* by cluster. **(E)** Expression of core EMT transcription factors and Vim by cell type.

Figure S7

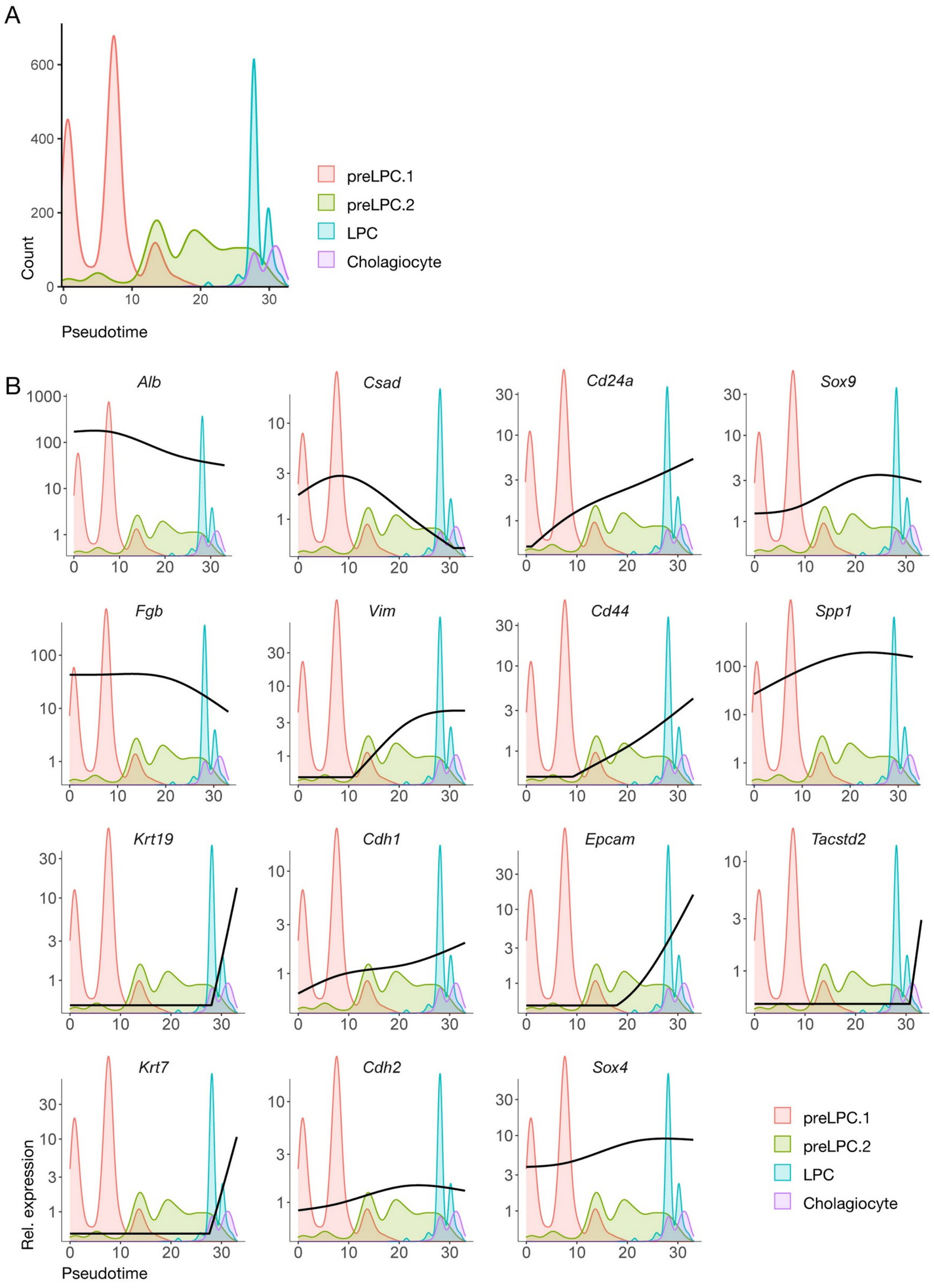
A
Count
Pseudotime
preLPC.1
preLPC.2
LPC
Cholagiocyte
B
Alb
Csad
Cd24a
Sox9
Fgb
Vim
Cd44
Spp1
Krt19
Cdh1
Epcam
Tacstd2
Krt7
Cdh2
Sox4
Rel. expression
Pseudotime
preLPC.1
preLPC.2
LPC
Cholagiocyte

**Figure S7. (A)** Distribution of cells during the LPC transition by cell type. **(B)** Expression curve of hepatic markers, cholangiocyte markers, and LPC markers along pseudotime.

Figure S8

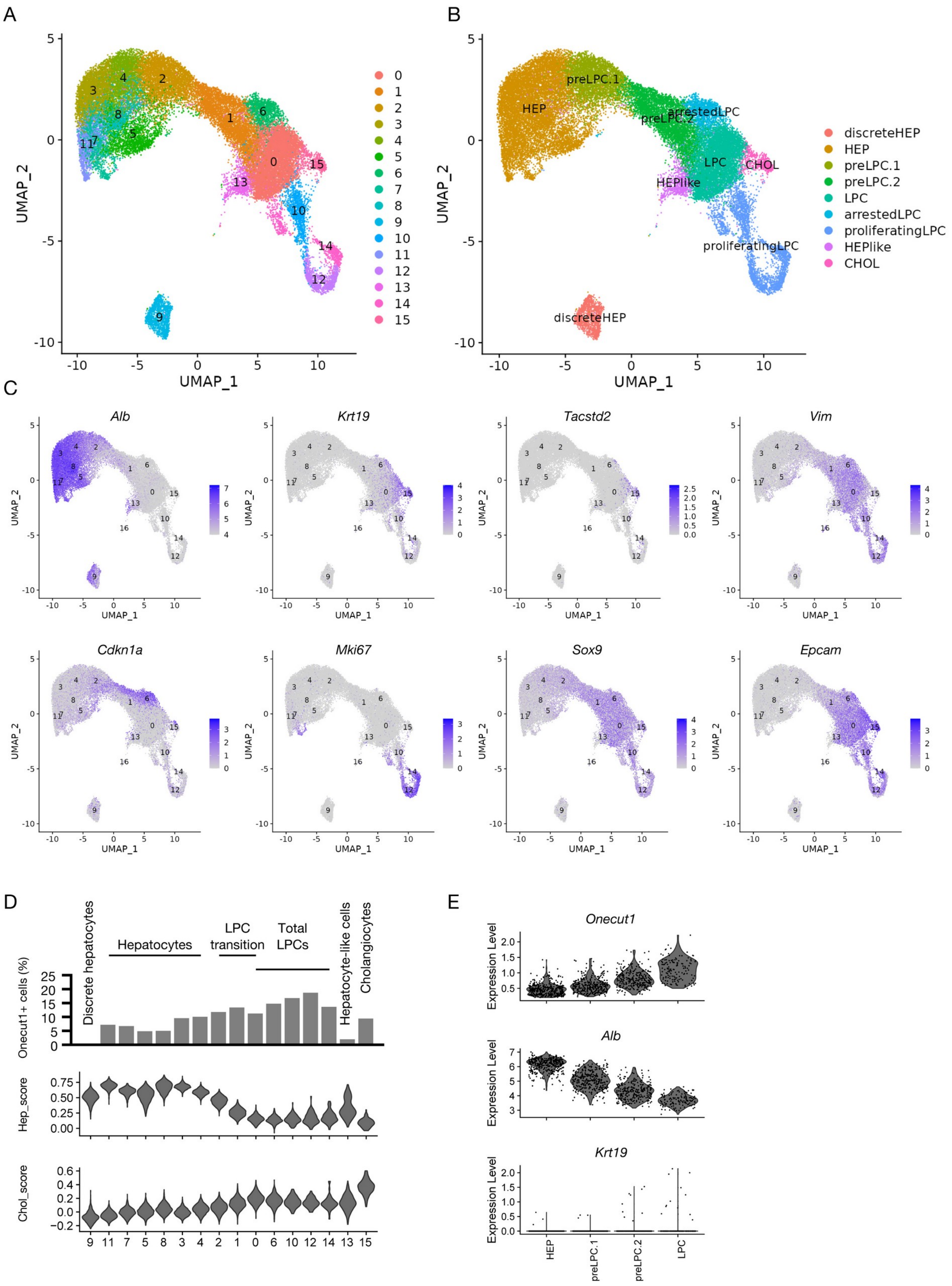
A
UMAP_1
UMAP_2
B
HEP
preLPC.1
preLPC.2
arrestedLPC
LPC
CHOL
HEPlike
proliferatingLPC
discreteHEP
C
Alb
Krt19
Tacstd2
Vim
Cdkn1a
Mki67
Sox9
Epcam
D
Discrete hepatocytes
Hepatocytes
LPC transition
Total LPCs
Hepatocyte-like cells
Cholangiocytes
Onecut1+ cells (%)
Hep_score
Chol_score
E
Onecut1
Alb
Krt19
Expression Level
HEP
preLPC.1
preLPC.2
LPC

**Figure S8. Analyses on sorted YFP$^+$ cells. (A)** Single-cell RNA-seq profiling of hepatocyte-derived cells by cluster. **(B)** Hepatocyte-to-cholangiocyte transdifferentiation continuum by cell type. Mature cholangiocytes were defined as a *Krt19*$^+$*Tacstd2*$^+$*Vim*$^-$ cluster. **(C)** Expression of the indicated genes in hepatocyte-derived cells. **(D)** Accumulation of *Onecut1*$^+$ cells with hep_score and chol_score values in control mice by cluster. **(E)** Expression of *Onecut1* in *Onecut1*$^+$ cells extracted from enriched clusters by cell type. Cholangiocytes were omitted due to small sample size.

Figure S9

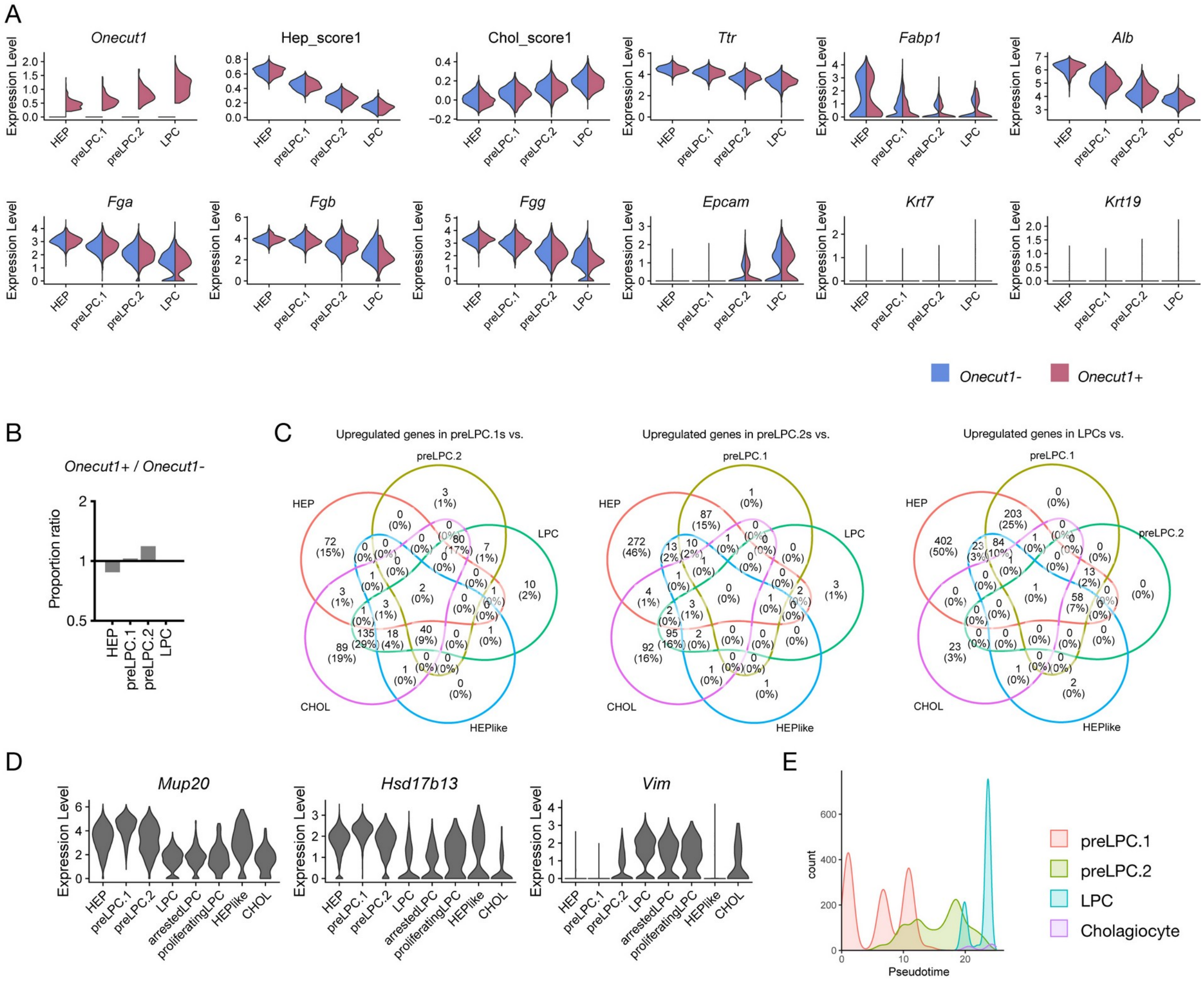

**Figure S9. Analyses on sorted YFP⁺ cells. (A)** Expression of the indicated genes in *Onecut1*⁻ and *Onecut1*⁺ cells by cell type. **(B)** Ratios of proportions of components in *Onecut1*⁺ cells to those in *Onecut1*⁻ cells by cell type. **(C)** Venn diagram of upregulated genes in preLPC.1s, preLPC.2s and LPCs. **(D)** Expression of common upregulated genes extracted in (B) by cell type. **(E)** Distribution of cells during the LPC transition by cell type.

Figure S10

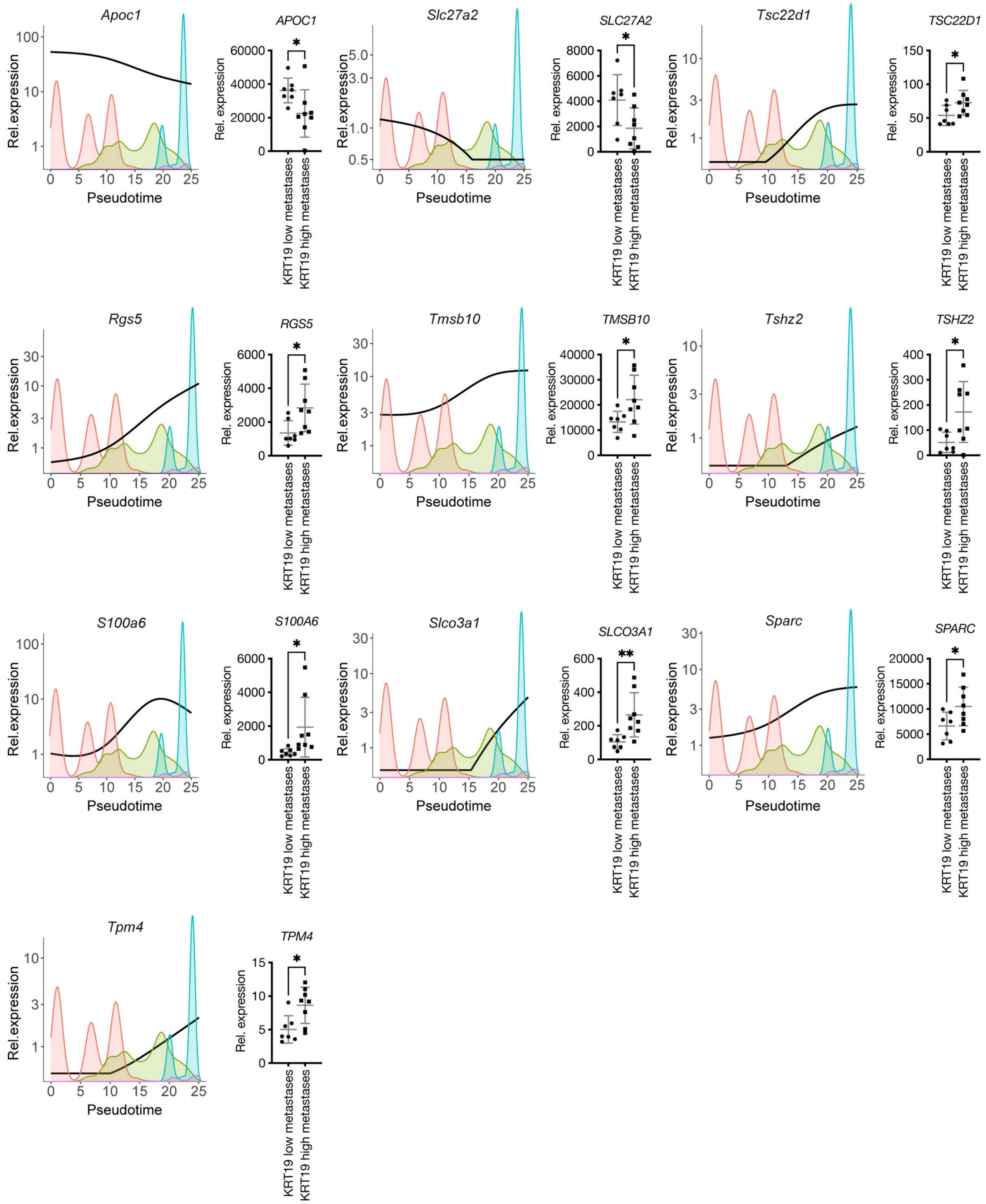
Apoc1
APOC1
Slc27a2
SLC27A2
Tsc22d1
TSC22D1
Rgs5
RGS5
Tmsb10
TMSB10
Tshz2
TSHZ2
S100a6
S100A6
Slco3a1
SLCO3A1
Sparc
SPARC
Tpm4
TPM4
Rel.expression
Rel. expression
Pseudotime
KRT19 low metastases
KRT19 high metastases
*
**

**Figure S10. Analyses on sorted YFP$^+$ cells.** Expression of common DEGs shown as an expression curve along pseudotime (left) or by expression in human HCC data (right). * indicates a significant difference by a t test. The expression of all genes upregulated in *KRT19*$^{high}$ metastases also increased along transdifferentiation pseudotime, while the expression of all genes downregulated in *KRT19*$^{high}$ metastases decreased along transdifferentiation pseudotime.

Figure S11

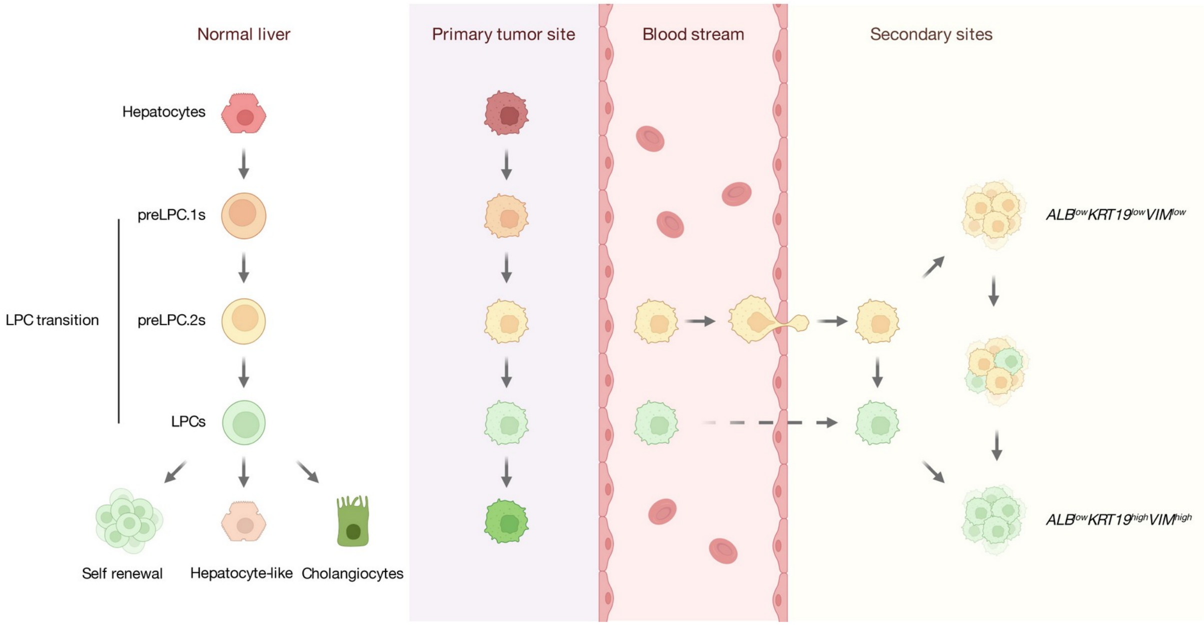


**Figure S11**. Scheme of the LPC transition during hepatocyte-to-cholangiocyte transdifferentiation, with possible acquisition of metastatic potential.
